# Efficient Delivering of a Photodynamic Therapy Drug into Cellular Membranes Rationalized by Molecular Dynamics

**DOI:** 10.1101/2023.10.16.562486

**Authors:** Basak Koca-Findik, Ilya Yakavets, Henri-Pierre Lassalle, Saron Catak, Antonio Monari

## Abstract

Photodynamic Therapy (PDT) represents a most attractive therapeutic strategy to reduce side effects of chemotherapy and improve the global quality of life of patients. Yet, many PDT drugs suffer from a poor bioavailability and cellular intake, and thus, drug delivery strategy are mandatory. In this article we rationalize the behavior of a temoporfin-based PDT drug, commercialized under the name of Foscan, complexed by two β-cyclodextrin units, acting as drug carriers, in presence of a lipid bilayer. Our all atom simulations have unequivocally shown the internalization of the drug-deliovering complex and suggest its possible spontaneous dissociation in the lipid bilayer core. The factors favoring penetration and dissociation have also been analyzed, together with the membrane perturbation due to the interaction with the drug carrier complex. Our results confirm the suitability of this encapsulation strategy for PDT, and rationalize the experimental results concerning its efficacy.

## Introduction

Photodynamic Therapy (PDT) is a promising therapeutic approach,[1–5] which combines the cytotoxic efficiency of a given drug with its activation by suitable electromagnetic radiation to improve the spatial and temporal selectivity of the treatment, and hence minimize side-effects. These features make PDT particularly promising in the field of cancer treatments [2,6], allowing to bypass the usual systemic toxicity of conventional chemotherapeutic agents, such as cis-platin [7] and its derivatives [8], which despite their efficiency induce important burdens on the quality of life of the patients. Furthermore, conventional chemotherapy agents may also lead to strong resistance effects [9,10], which result in a significant reduction of their efficiency and clinical applicability. From a fundamental point of view PDT relies on the administration of a drug which is inert in the dark, and thus of low toxicity. The excitation by suitable wavelengths induces the population of the drug’s excited states manifold which may favor the production of reactive oxygen species (ROS). The latter will further attack biological macromolecular structures, such as lipid membranes, proteins, or nucleic acids, and, thus, lead to cell death [11]. One of the most common photophysical pathway exploited in PDT relies on the activation of molecular oxygen to its singlet excited state ^1^O_2_, which has the capacity to easily oxidize biological molecules. The activation of molecular oxygen to ^1^O_2_ requires the population of the drug’s triplet states via intersystem crossing (ISC), whose facility and time-scale is therefore crucial in dictating the global therapeutic efficiency. Other possible PDT strategies are based on the production of superoxide or peroxy-radicals, usually achieved through excited-state electron transfer [12]. Porphyrins and phthalocyanines are probably among the most common PDT agents exploited nowadays both in clinical applications and in fundamental research approaches [5,13,14]. This is mainly due to their high extinction coefficients and their facile ISC, which, furthermore, can be modulated by the suitable inclusion of heavy metal atoms, such as Pt, Zn, or Pd, in the coordination sphere constituted by the porphyrin core [15,16].

Despite the clear success of PDT in cancer treatment, especially for esoph
ageal, uterine and mouth cancers, as well as in the treatment of different viral [17], bacterial [18], and auto-immune diseases [19], such as psoriasis [20], PDT still suffers from some critical drawbacks, which should be properly considered [21].

A PDT drug should ideally absorb in the red and infrared portion of the electromagnetic spectrum, and specifically in the therapeutic window, i.e. at wavelengths biological tissues are transparent. This prerequisite is actually crucial to assure the highest penetration of the activating light, and hence the treatment of deeper lesions. Furthermore, the accumulation of the drug into cancer cells is also suitable to enhance the therapeutic efficiency of PDT. For this reasons, specific strategies based on the functionalization of PDT drug cores with specific antibodies, targeting receptors which are over expressed on cancer cells, have been proposed [22–24].

Nevertheless, an additional limitation occurs more specifically for porphyrin-based PDT agents, which due to their extended hydrophobic π-conjugated system are prone to aggregation, thus strongly limiting their bioavailability and their possible oral or intravenous administration. Furthermore, aggregation may also significantly alter the photophysical properties of the drug, shifting the absorption spectrum maxima, and more importantly opening possible competitive pathways, such as vibrational relaxation, impeding an efficient ISC, and thus ^1^O_2_ activation.

As a matter of fact, a chlorin-based temoporfin (m-THCP, Figure 1) [25–27], commercialized under the name of Foscan, is recognized as an efficient PDT agent, especially for the treatment of squamous cell head-and-neck cancer [28,29], even in advanced stages and for palliative purposes [30]. Different studies, including pre-clinical and clinical tests, are also devoted to its possible use to treat other cancer types, including, for instance, ovarian cancer [31]. Furthermore, the behavior of m-THCP and chlorin has also been studied computationally, including its embedding in liposomes by Nogueira group [32]. Despite its success, m-THCP strongly suffers from aggregation issues, which severely limit its bioavailability [33–36].

**Figure 1.**
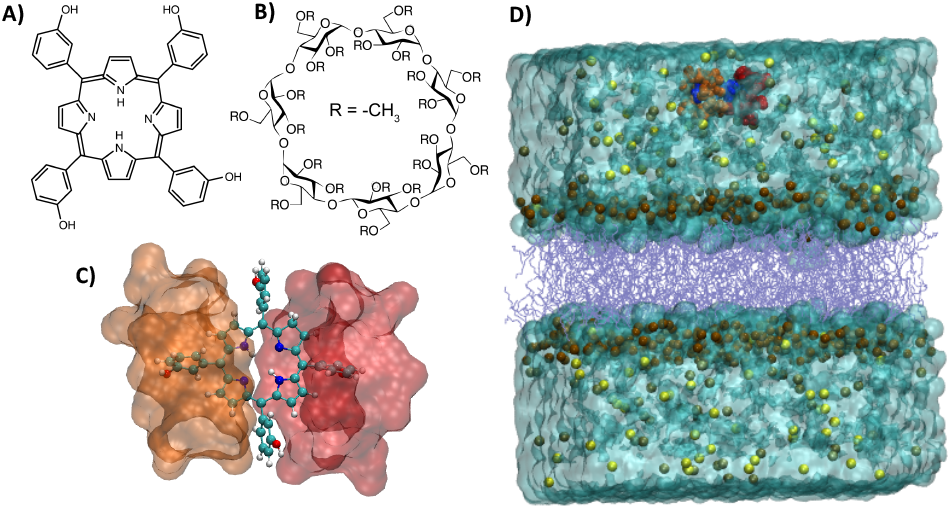
Chemical formula of m-THPC (A) and BCD (B). C) The 1:2 m-THPC/BCD complex in which m-THPC is represented with ball and sticks while BCD as transparent surfaces. D) Illustration of the whole simulation box including the lipid bilayer, water bulk, the m-THPC/BCD complex and K^+^, Cl^-^ ions.

Recently, by using both molecular modeling and simulations and experimental approaches [37–39], we have shown that m-THPC is able to persistently interact with lipid bilayers, mimicking cellular or mitochondrial membranes. This leads to favorable conditions for PDT, since the drug can activate ^1^O_2_ close to the potentially reactive target, i.e. the oxidizable lipid double bonds. We have also shown that m-THPC can be easily and persistently encapsulated in a 1:2 ratio by β-cyclodextrin derivatives [37,38]. The latter represents a strategy which could limit aggregation and, thus, strongly enhance bioavailability. By using spectroscopic titration, the thermodynamic favorable encapsulation process has also been characterized in terms of its binding free energy [38]. We have also shown that the encapsulation takes place by an asynchronous mechanism involving the sequential inclusion of the two cyclodextrin units [38]. On the other hands, we have confirmed that encapsulation is not altering the optical and photophysical properties of m-THPC, thus maintaining its ideal properties for PDT applications [37]. This observation is also confirmed by the fact that cyclodextrin-encapsulated m-THPC is highly cytotoxic or toxic under irradiation [40–42].

Yet, the precise mechanisms through which encapsulated m-THPC is interacting with biological systems, such as cellular membranes, has not been characterized yet. In particular, the mechanisms leading to the internalization of the encapsulated drug and its release in the proximity of the cellular membrane remain elusive. Furthermore, the accessibility of the encapsulated chromophore to molecular oxygen, which is a crucial parameter to allow ^1^O_2_ activation and, thus, efficient PDT applications, should be better rationalized. For these reasons in this contribution we perform molecular dynamic simulations of β - cyclodextrin-encapsulated m-THPC in presence of a model lipid bilayer mimicking a cellular membrane, exceeding the μs time-scale. Our results unambiguously show that the drug delivering complexes is easily internalized into lipid membranes. Moreover, its internalization may also correlate with its spontaneous dissociation and to the release of the PDT drug in the lipid membrane, which is coherent with the experimentally observed cytotoxicity under irradiation.

## Computational Methodology

We have performed long-scale all atom molecular dynamic (MD) simulations exceeding the μs time-scale in three replicas of a β-cyclodextrin (BCD) encapsulated m-THPC interacting with a phosphatidylcholine (POPC) lipid bilayer surrounded by a water buffer and physiological salt concentration (Figure 1). The lipids have been modeled using the lipid14 force field [43]. In the case of cyclodextrin and m-THPC, the force fields are the same as those used in our previous contributions [37], which are based on Glycam_06 [44] and Generalized Amber Force field (GAFF) [45], respectively. Water is instead modeled using the TIP3P force field [46]. All the simulations have been performed using the NAMD code [47,48] and visualized and analyzed with VMD [49]. The initial systems have been built using the Charmm-Gui web interface [50]. In addition, the encapsulated m-THPC was also simulated in a water box. In the case of the complex in bulk water a cubic simulation box having an initial size of 100.0 x 54.1 x 54.6 Å^3^ was selected. The complex interacting with the bilayer system was placed in a 96.0 x 91.0 x 120.0 Å^3^ box. Each membrane leaflet was composed of 150 POPC units and has been hydrated by 14808 water molecules.

All simulations have been performed in the constant pressure and temperature ensemble (NPT) at 1 atm and 300K, which have been enforced using the Langevin barostat and thermostat, respectively. Electrostatic interactions have been obtained using the Particle Mesh Ewald (PME) summation considering a cut-off of 9 Å. For each simulation, hydrogen mass repartition (HMR) [51] was applied to slow the highest frequency vibrations involving hydrogens, thus allowing the use of a 4.0 fs time step to integrate the Newton’s equations of motion. After 60000 conjugated gradient minimization steps, the initial systems have been thermalized and equilibrated by gradually removing the constraints on the heavy atoms of the membrane. This has been performed in three consecutive steps of 36 ns each. The stability of the systems has been checked by analyzing the time evolution of the root mean square deviation analysis. To better characterize the internalization of the complex and its partial dissociation the linear interaction energy (LIE) time-series has been obtained using the Amber utilities [52–54]. Topologies and trajectories for the three replicas are publicly available on a Zenodo repository (https://zenodo.org/records/11530985).

## Results and Discussion

Coherently with our previous work, the 1:2 m-THPC/BCD complex is highly stable in the water bulk (see ESI), confirming a favorable encapsulation. The behavior of the same m-THPC/BCD complex in presence of a lipid bilayer can be appreciated from the snapshots collected in Figure 2, which are relative to the first replica. The drug delivery complex, which was initially placed in the water bulk, is rapidly developing favorable interactions with the lipid bilayer. After about 15 ns, the complex reaches the polar head region and strengthen the interaction network at the water/lipid interface. This can also be appreciated by the distortion of the membrane surface, which forms a convex area in the vicinity of the m-THPC/BCD complex.

**Figure 2.**
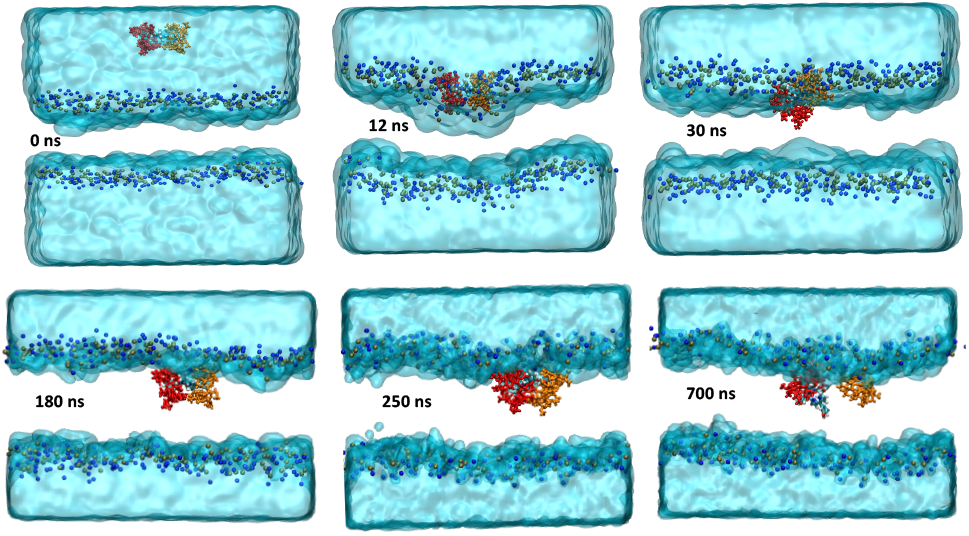
Snapshots extracted along the MD simulation showing the internalization of the m-THPC/BCD complex and its spontaneous dissociation in the lipid bilayer.

However, this situation is evolving quite rapidly, and after about 30 ns the complex is buried deeper in the polar head region. Interestingly, in this state the deformation of the lipid membrane remains pronounced. Rather surprisingly, in the 180-250 ns range we observe that the m-THPC/BCD complex is actually spontaneously penetrating inside the lipid bilayer without experiencing any dissociation of its constitutive units. Interestingly, the spontaneous penetration is also confirmed by all the replicas, with similar time-scales spanning the hundreds of ns regime, as reported in ESI. This behavior was unexpected, especially considering the relatively large size of the complex and the rather hydrophilic propensities of cyclodextrin. Yet, the passive internalization of the drug-delivering unit should be regarded as highly beneficial for PDT, and in general for pharmacological applications.

After this step, the fully internalized complex still resides in the vicinity of the polar head region, experiencing some important rotational flexibility, most probably to maximize the interactions between the lipid polar groups and the cyclodextrin hydrophilic moieties. However, in the case of the first replica, after about 450 ns a major reorganization may be observed. Indeed, the strong interactions of the BCD units with the polar heads actually induce the opening of the cyclodextrin complex, its partial dissociation and, thus, the exposure of the m-THPC core to the hydrophobic lipid environment. This behavior can also be appreciated by analyzing the time evolution of the distance between the center of mass of m-THPC and the BCD units (Figure 3A) which clearly shows a first partial release of one BCD. Indeed, the distance between one BCD moiety and m-THPC increases firstly to around 10 Å from 200 to 500 ns, indicating the partial opening of the complex. Afterwards, a sharp discontinuity in the time series can be observed indicating the complete release of the BCD moiety, as confirmed by the distance overcoming the 18-20 Å range.

**Figure 3.**
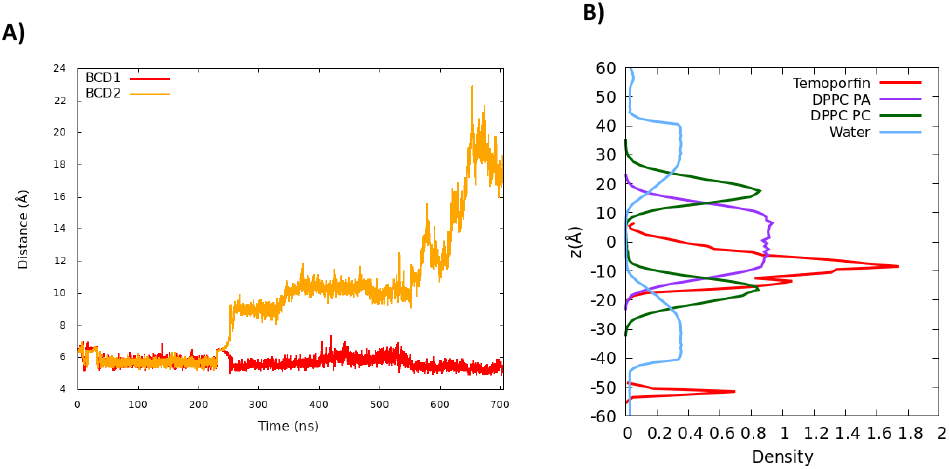
A) Time series of the distance between the centers of mass of m-THPC and the two BCD units. B) Density profile along the membrane axis of m-THPC, BCD, and lipid polar heads. The secondary peak of m-THPC is representative of the pre-equilibrated situation in which the complex is in the water bulk. This Figure refers to the first replica the results for the other two are reported in ESI.

The time scale of our MD simulation is not sufficient to enforce the spontaneous release of the second BCD unit, yet we may hypothesize that this process should take place given the affinity of the partially exposed m-THPC for hydrophobic environments. On the other hand, even the partial opening of the m-THPC/BCD complex is actually sufficient to allow the encounter between the chromophore and membrane-dissolved molecular oxygen, which is necessary for energy transfer and hence ^1^O_2_ activation. Interestingly, as shown from the density profile along the membrane axis (Figure 3B), m-THPC is residing close to the reactive lipid double-bonds, partially overlapping with their distribution, hence assuring the production of singlet oxygen in close proximity to its chemical target, and, consequently, decreasing the possibility of quenching.

As shown in ESI the spontaneous partial opening of the drug delivery complex upon internalization is also observed in the case of replica 3 with a time scale of about 600 ns. On the other hand, no dissociation has been observed in the case of replica 2 after 1 μs. This different behavior may suggest the coexistence of dissociated and associated complexes or to the necessity to sample longer timescales. Yet, the observation of the complex opening on two over three replicas clearly underlines its potential possible occurrence. Furthermore, as shown in ESI, we may observe that in case of replica 2, for which the spontaneous opening of the complex does not take place, the root mean square (RMSD) of the lipid membrane is not plateauing, suggesting that the large size of the complex requires more pronounced conformational reorganization of the lipid environment. On the contrary, a clear stability of the RMSD time evolution can be observed in the case of the replicas experiencing dissociation.

To better analyze the interplay between the complex internalization and the membrane stability, in Figure 4 we also report the time evolution of some membrane structural parameters, obtained using the MEMBPLUGIN extension of VMD [55], namely the area per lipid and the membrane thickness. Note that for comparison in ESI we also report the area per lipid obtained for a simulation of the same membrane in absence of the PDT drug-delivering complex. In Figure 4 only the results relative to replica 1 have been reported, while the behavior of the other replicas can be appreciated in ESI. Interestingly, and despite the local strong perturbation, we may observe only limited deviation from ideality for the two descriptors. The membrane thickness is slightly decreasing compared to the initial situation representative of an equilibrated bilayer and rapidly stabilizes at around 38 Å. On the other hand, the area per lipid is initially decreasing, corresponding to the formation of the pre internalized complex which is accompanied by a local strong deformation of the membrane. After the internalization, as can be expected, the area per lipid slightly increases by 3-4 Å^2^ with respect to an ideal membrane, and despite some oscillations it remains globally stable all along the MD simulation.

**Figure 4.**
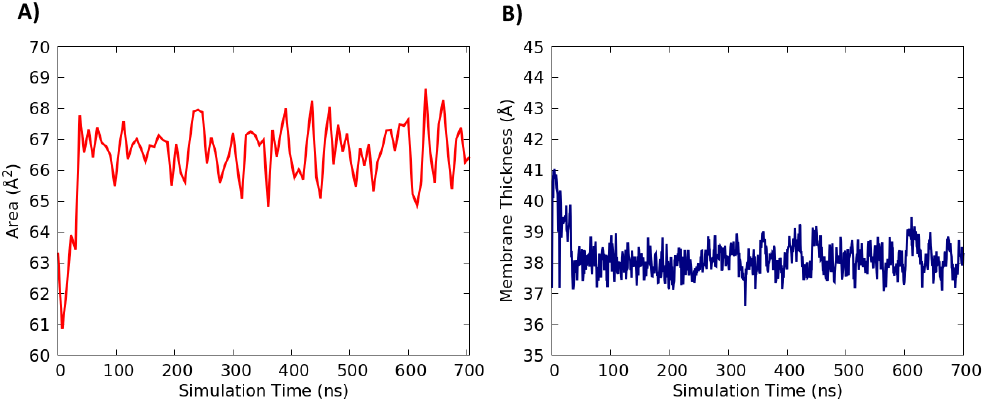
Time evolution of the area per lipid (Å) and membrane thickness (B) along the MD simulation for the first replica.

These data confirm that the internalization and the dissociation of the m-THPC complex are not accompanied by a strong perturbation of the bilayer. This aspect should also contribute to keep the drug toxicity in the dark at a relatively modest level, further confirming its suitability for PDT applications. A similar behavior can be observed for the structural parameters of replica 3 which leads to the complex partial dissociation, while, coherently with the time evolution of the lipid RMSD, replica 2, involving the internalized but undissociated complex, shows a more pronounced deviation from ideality.

From a molecular ant atomistic point of view, the main interactions leading to both the complex internalization and its eventual dissociation mainly involve the polar head and the peripheral substituents of the BCD complex as highlighted in Figure 5 for the first replica.

**Figure 5.**
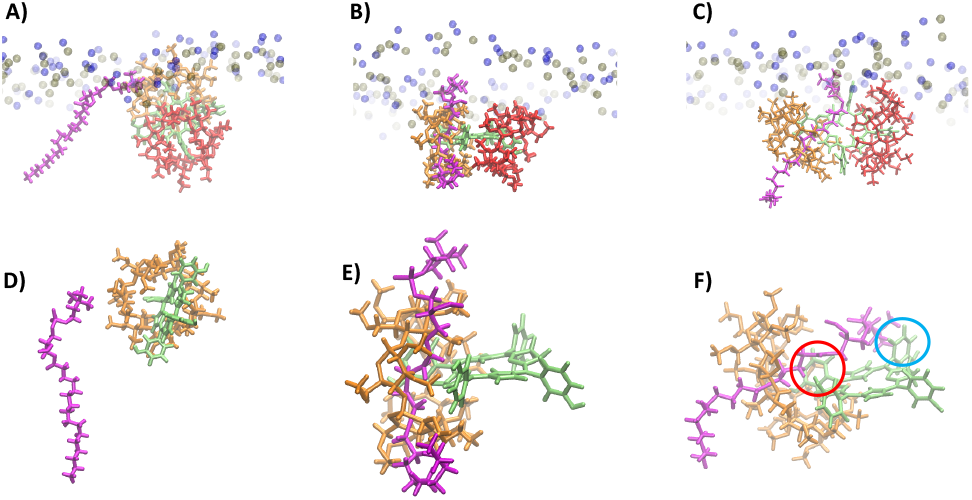
Main interactions developed by the m-THPC/BCD complex and the lipid polar heads leading to the complex partial dissociation. A) interaction between BCD and the polar head of one lipid chain leading to recognition, B) insertion of the lipid tail in the interface region between the BCD units destabilizing the complex, C) stabilization of the exposed m-THPC by interactions with the lipid tail and polar head. Zooms of the previous states are also provided in panel D (recognition), E (complex destabilization), and F (m-THPC stabilization). Note that in panels D-F only the opening BCD is represented. The formation of cation-π and π-alkyl interactions are highlighted in panel F by red and blue circles, respectively.

Indeed, we may observe that the dissociation proceeds through a subtle balance of hydrophobic and hydrophilic interactions involving the BCD complex and the nearby lipids. Specifically, we may recognize an initial step in which the lipid polar head, and particularly the N(CH_3_)^+^ moiety interacts with the methoxy oxygens of the cyclodextrin, thus driving the recognition and the formation of a short-lived state (Figure 5A, D). This intermediate state is further evolving, since the hydrophobic tail of the same lipid develops strong interactions with the BCD core, which result in its insertion at the interface between the two cyclodextrin units (Figure 5B, E).

Afterwards, and following the partial dissociation of the complex, m-THPC is stabilized by diverse factors involving cation-π interactions between the lipid polar heads and the peripheral phenyl rings, and π-alkyl interactions exerted by the hydrophobic tail (Figure 5C, F). Thus, the presence of the lipid tails and globally the amphiphilic character of the lipids appear as fundamental to disrupt the interaction network maintaining the BCD/m-THPC complex. Furthermore, the subsequent stabilization of the exposed m-THPC and the newly formed interaction network may be deemed responsible to hamper the subsequent reformation of the complex, making its opening irreversible. As concerns the second BCD unit its methoxy oxygen atoms also form favorable interactions with the polar head, contributing to maintain this unit firmly anchored at the interface region. A similar pattern, involving again the subtle balance between hydrophobic and hydrophilic interactions, is also observed for the third replica, which results in the partial dissociation of the complex, as shown in ESI. This further confirms the relevance of the observed dissociation.

These effects can also be appreciated by the analysis of the time evolution of the LIE as reported in Figure 6 for the first replica and in SI for all the others. Indeed, we can observe that the rapid internalization of the complex involves a fast decrease of both the electrostatic and van der Waals interactions terms with all the components. The opening of the complex is instead characterized by a very sharp increase in LIE, which is correlated with the instability brought by the approaching of the lipid tail as shown in Figure 5. After the partial opening of the complex we also observe, as expected, that the interaction between m-THPC and the leaving BCD unit (Figure 6D) is strongly decreased compared to the remaining group (Figure 6C). While the difference appears less significant the same reduction of the interaction strengths between the leaving BCD and the membrane polar head can be observed. Similar effects can also be sketched from the other replicas, notably concerning the strong stabilization upon the complex internalization. However, especially for replica 3, since the complex is only partially dissociating the interaction energies are much more conserved all along the MD simulation.

**Figure 6.**
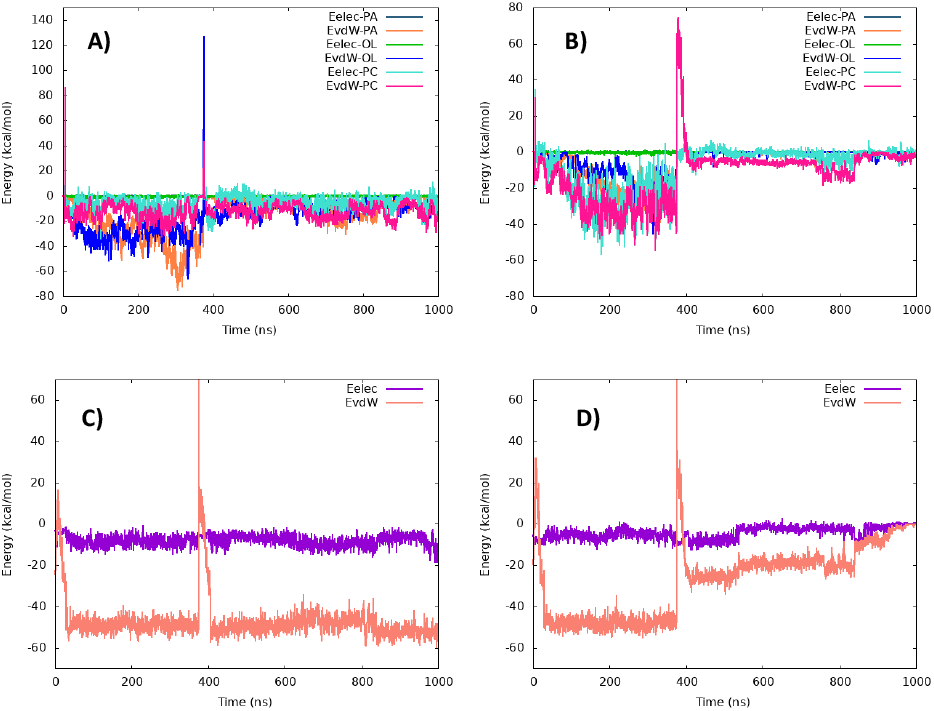
Time evolution of the LIE calculated for the first replica between the lipid moieties and the first (A) and second (B) BCD unit, respectively. The time evolution of the LIE between m-THPC and thie first C) and second (D) BCD unit is also reported.

## Conclusions

By using long-scale all atom MD simulation we have shed light on the drug delivering propensity of a cyclodextrin-complexed temoporfin agent, which is used in clinical PDT application. In particular, and rather surprisingly, we have shown that the drug-delivering complex not only interacts with the lipid polar head but also leads to its spontaneous internalization in the hundreds of ns time-scale. Even more surprisingly, we have shown that the internalized complex, may experience partial dissociation in the μs time-scale, at least for two over three replicas. We have shown that the partial dissociation is driven by the combined effects of the locking of one cyclodextrin at the polar head combined with the hydrophobic affinity of m-THPC for the lipid core. The whole process may happen in a sub-ms time-scale and, thus, should be biologically relevant, although a possible equilibrium between dissociated and undissociated complexes may be established. Furthermore, the combined effect of spontaneous internalization and dissociation is clearly beneficial and can justify the observed efficiency of the cyclodextrin-based vectorization in PDT. Indeed, while cyclodextrin is encapsulating and protecting the chromophore in water environment, notably avoiding unsuitable aggregation, it also readily targets temoporfin to the cellular membranes where it can exert its photo-cytotoxic effects.

Obviously, the accuracy of classical MD simulations relies heavily on the choice of the force field and its ability to reproduce satisfactorily the structural and dynamical properties of the system. The performance of lipid force fields in reproducing membrane properties, as well as passive internalization of small molecules has been recently reviewed [56,57], and despite the complexity of the systems and the interplay between different parameters and membrane conclusions, the outcomes are justifying the study of membrane penetration by force field based MD simulations.

While our study represents a first explanation at molecular and atomistic level of the subtle interplay leading to drug-delivering and vectorization in PDT applications, it also allows to better understand the molecular factors driving the interaction with lipid bilayer, which can be further optimized in a rational molecular design approach. In particular, the possibility to decorate the cyclodextrin with specific moieties, such as antibodies or peptides, specifically overexpressed in cancer cells is particularly attractive to enhance the selective deliver of the PDT agents, and will make the object of a further contribution.

## Supporting Information

RMSD of the lipid and ligands, time series of the most important distances describing the interaction leading to the BCD opening, description of the behavior of the different replicas and LIE time series. (file type, PDF).

## Acknowledgements

The authors thank GENCI and Explor as well as TUBITAK ULAKBIM High Performance and Grid Computing Center (TRUBA resources) and National Center for High Performance Computing of Turkey (UHeM) under grant number 1011062021 for computational resources. BKF and SC thank TUBITAK (Project Number: 120Z659) for financial support. BKF would also like to thank the French Embassy in Turkey for a joint Ph.D. grant. A.M. thanks ANR and CGI for their financial support of this work through Labex SEAM ANR 11 LABX 086, ANR 11 IDEX 05 02. The support of the IdEx “Université Paris 2019” ANR-18-IDEX-0001 and of the Platform P3MB is gratefully acknowledged.

## Entry for the Table of Contents

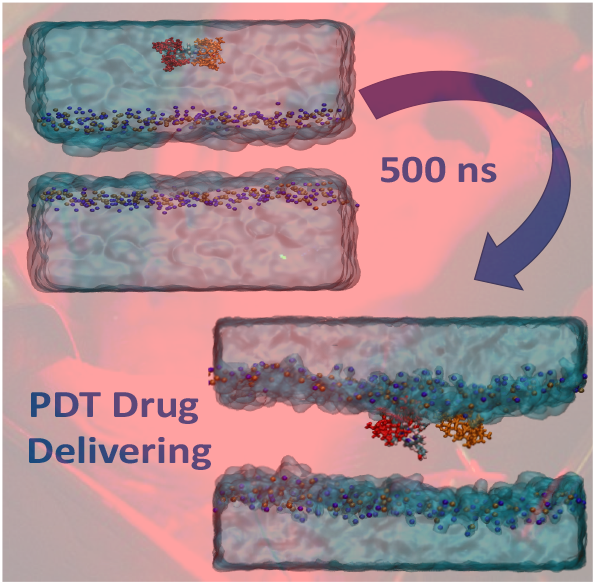

Institute and/or researcher Twitter usernames: @AntonioMonari

## Notes

Supporting information for this article is given via a link at the end of the document.

### Competing Interest Statement

The authors have declared no competing interest.

### Summary of Updates

Some replicas have been added and firther analysis of the MD simulations

