## Supplementary Information for "Efficient Delivering of a Photodynamic Therapy Drug into Cellular Membranes Rationalized by Molecular Dynamics"

a) Bogazici University, Department of Chemistry, Bebek 34342, Istanbul, Turkey. b) Université Paris Cité and CNRS, ITODYS, F-75006, Paris, France. c) Department of Chemistry, University of Toronto, 80 Saint George Street, Toronto, Ontario M5S 3H6, Canada. d) Université de Lorraine and CNRS, CRAN, UMR 7039, F-54000, Nancy, France. e) Université de Lorraine, Institut de Cancérologie de Lorraine, F-54000, Nancy.

AUTHOR INFORMATION

**Corresponding Author**

*

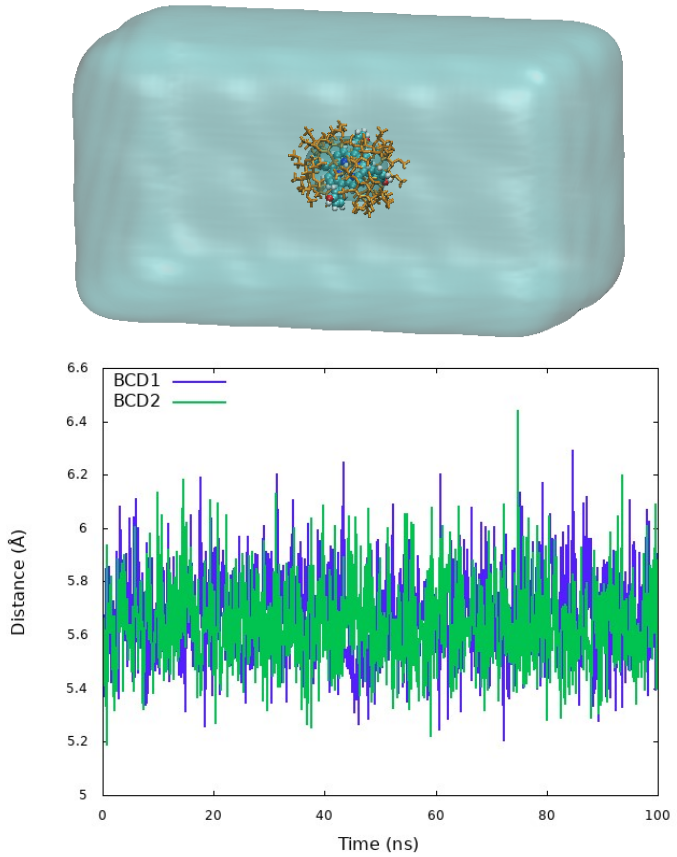


Figure S1. Equilibration MD during 100ns in NPT ensemble (100.0 x 54.1 x 54.6 Å$^3$) and time Evaluation of the interactions of BCD units with mTHPC in bulk water.

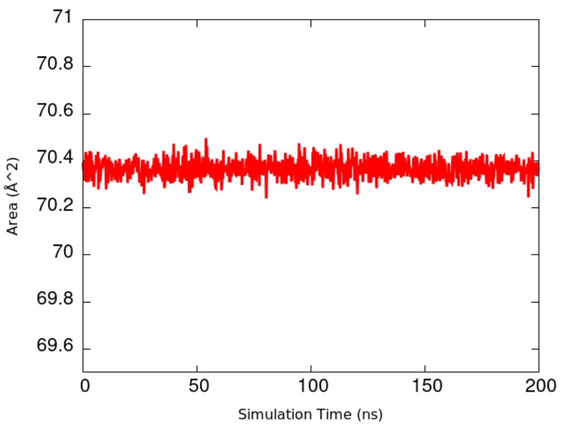


Figure S2. Area-per-lipid of POPC membrane used in simulations in the absence of the drug-delivering complex.

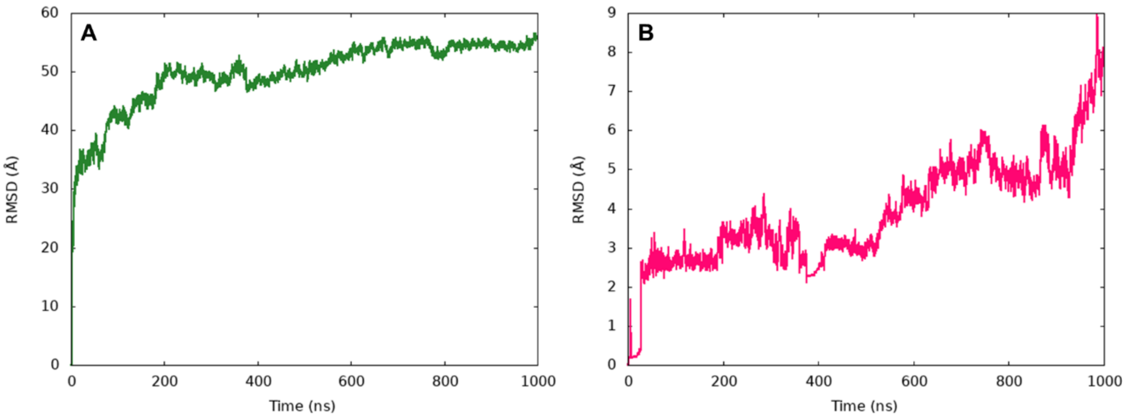


Figure S3. Root mean square deviation plots during 1μs equilibration MD of Replica-1 of the drug delivering complex in presence of the lipid bilayer. A) RMSD for the lipid and B. the complex. Note that the increase in the complex RMSD is due to the dissociation taking place.

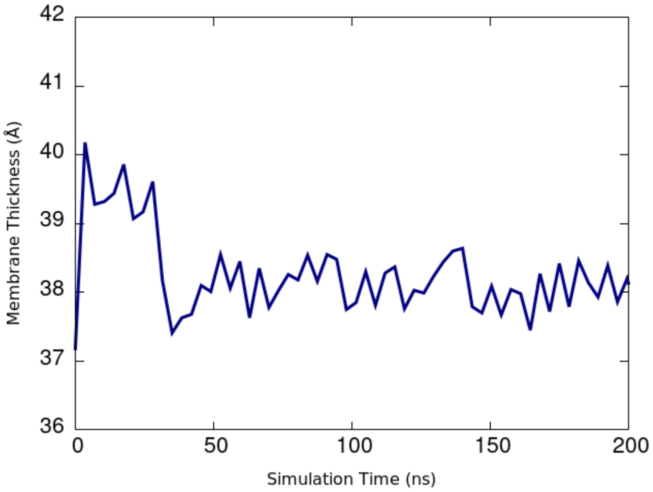


Figure S4. Time series for the membrane thickness of Replica-1 during the first 200ns corresponding to the internalization of the complex.

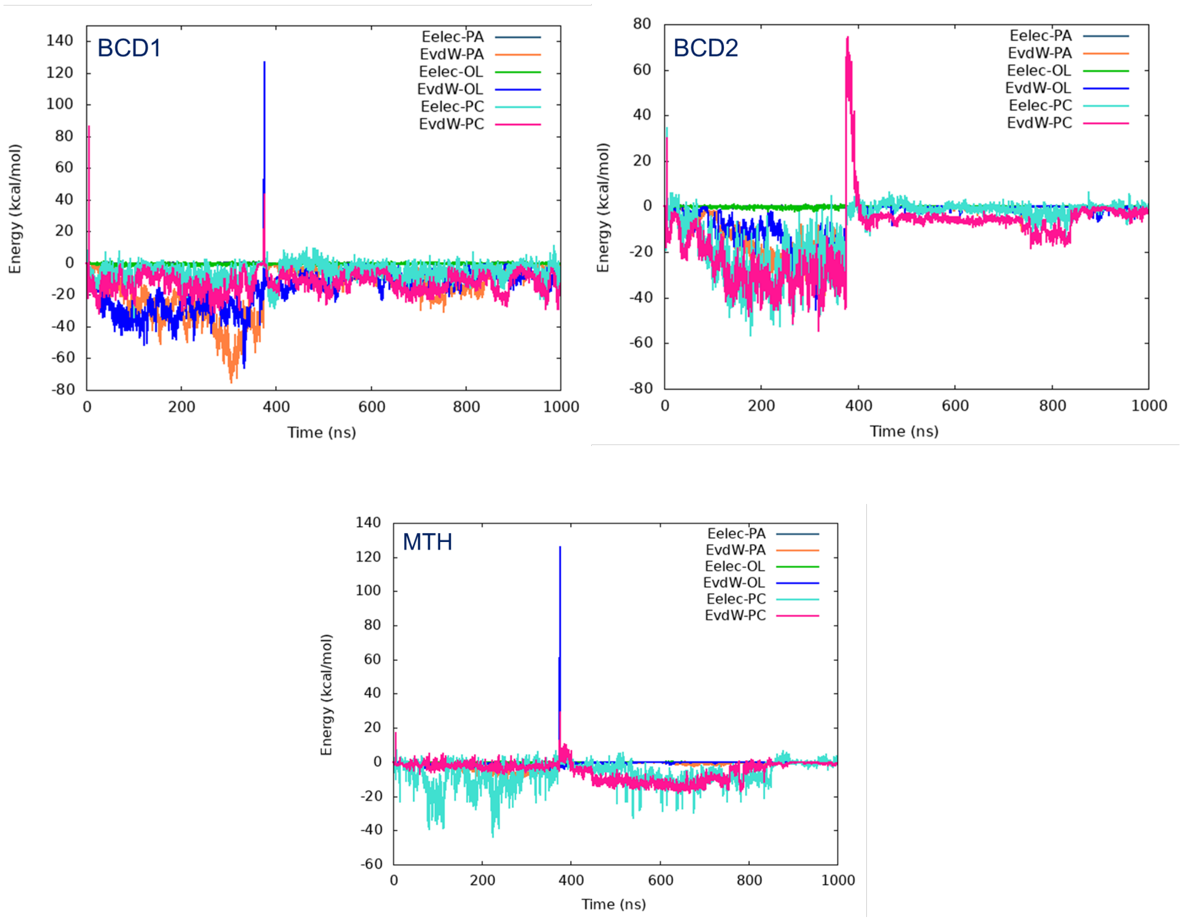


Figure S5. Time evolution of the LIE between the BCD and MTH and the lipid components for the first replica.

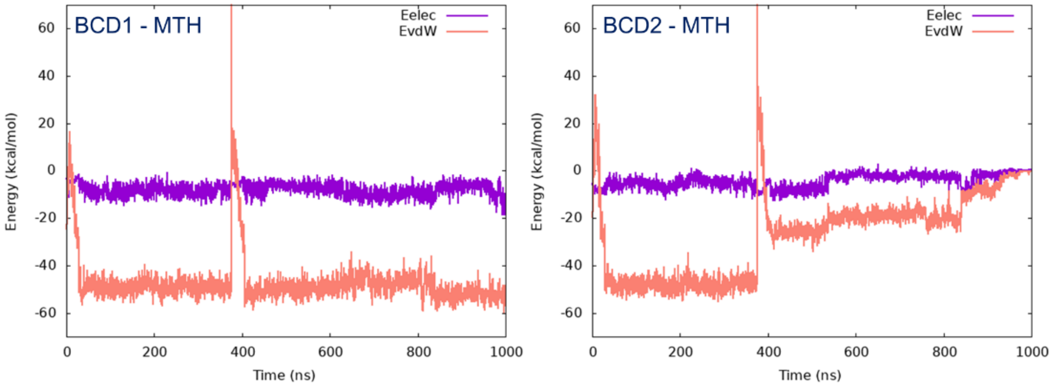


Figure S6. Time evolution of the LIE between the m-THPC and the BCD units for the first replica.

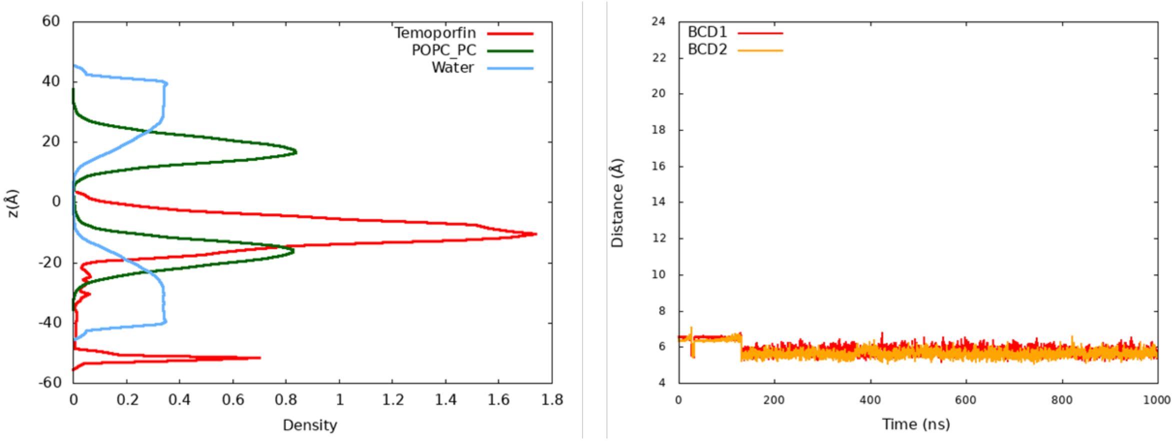


Figure S7. Density Profile (left) and time evaluation of the center of mass distances between each cyclodextrin units and mTHPC molecule (right) of Replica-2.

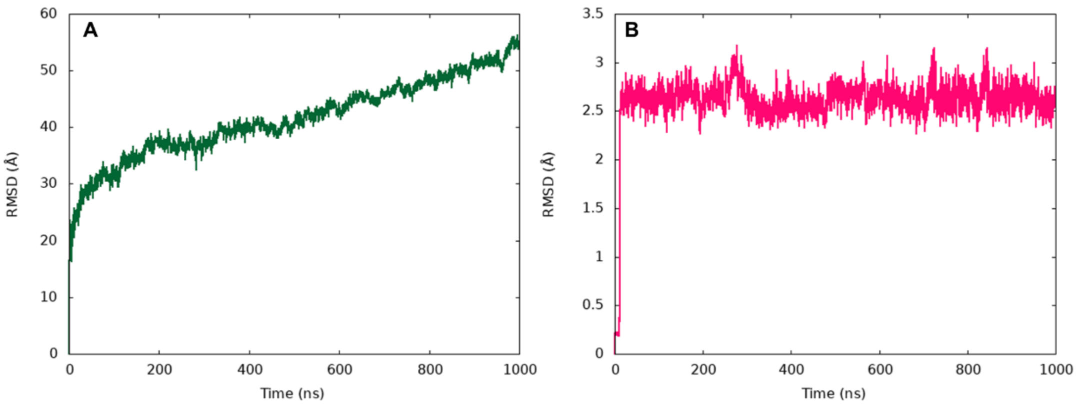


Figure S8. Root mean square deviation plots during 1µs equilibration MD of Replica-1 of the drug delivering complex in presence of the lipid bilayer. A) RMSD for the lipid and B. the complex. The increase in the membrane RMSD is most probably caused by the larger perturbation caused by the undissociated complex.

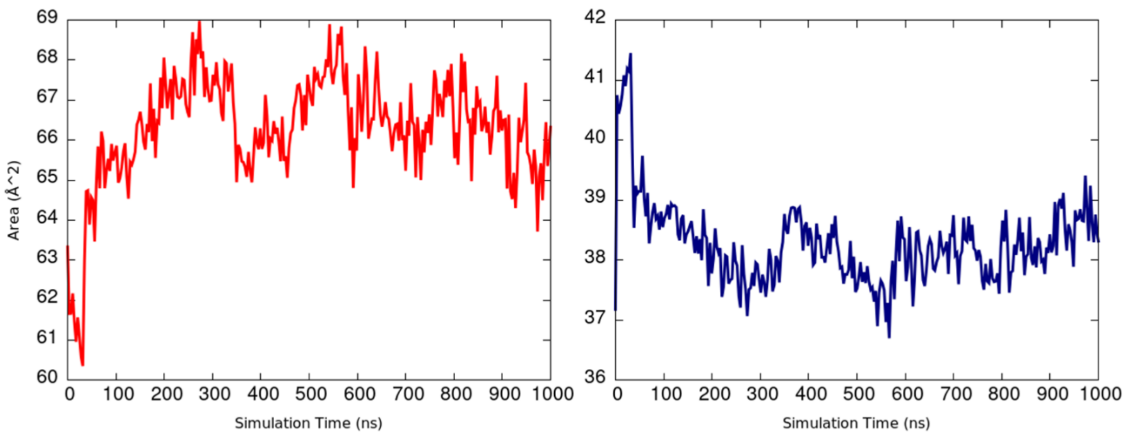


Figure S9. Time series of the area per lipid (red, left) and membrane thickness (navy blue, right) of Replica-2

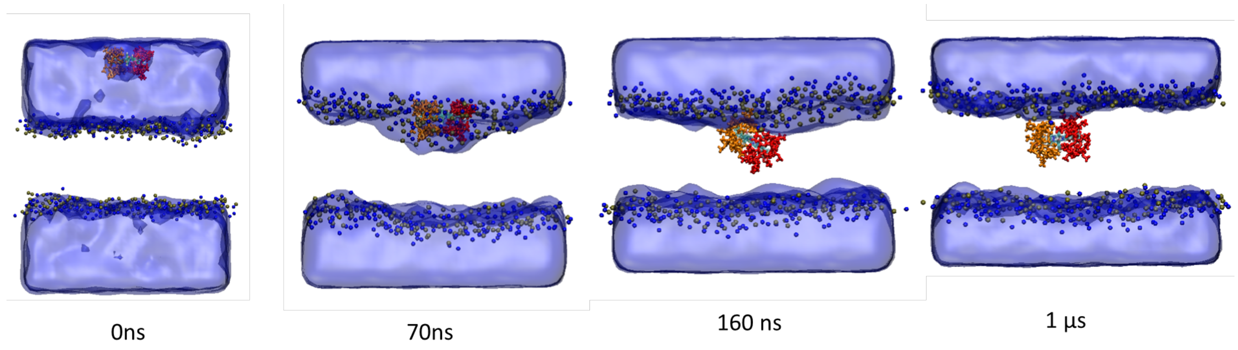


Figure S10. Snapshots representing the complex internalization from the replica-2 simulation.

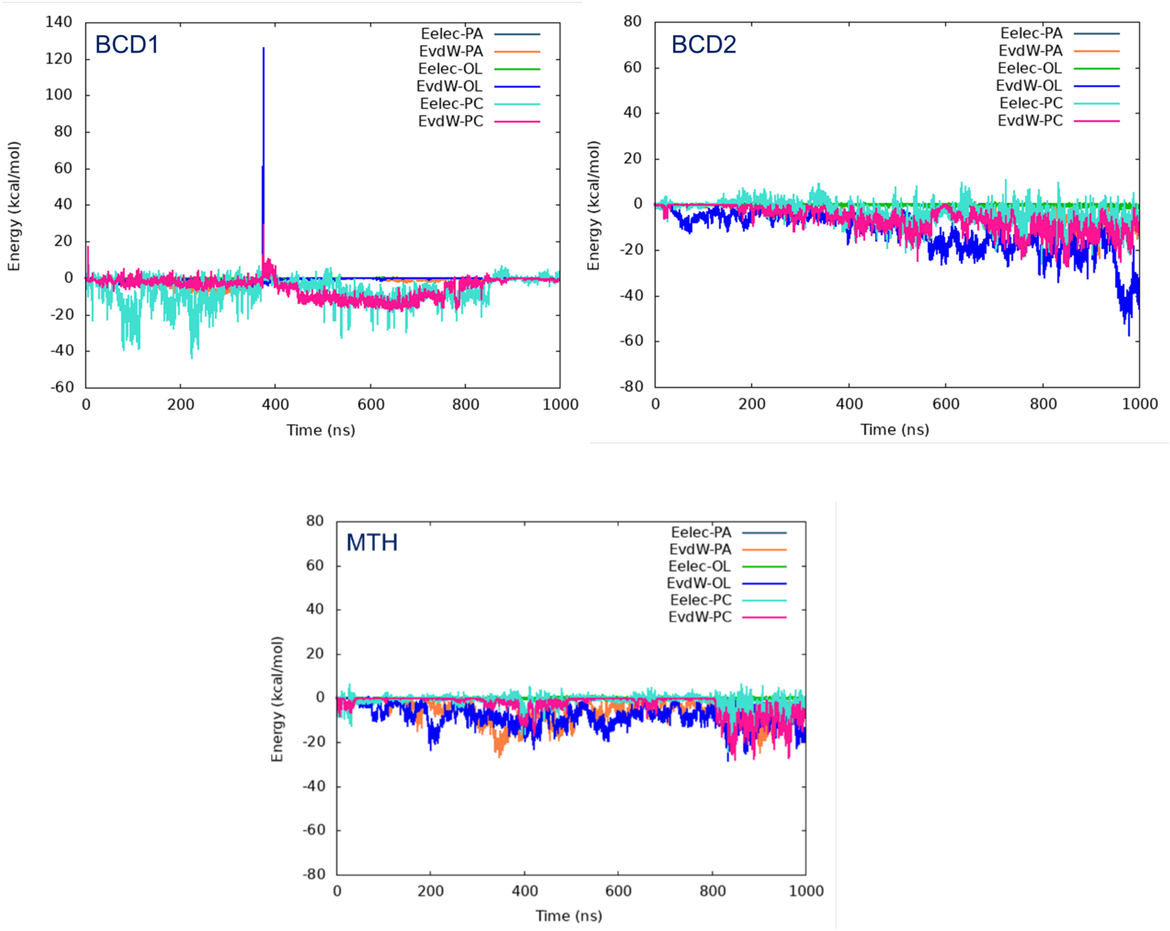


Figure S11. Time evolution of the LIE between the BCD and MTH and the lipid components for the second replica.

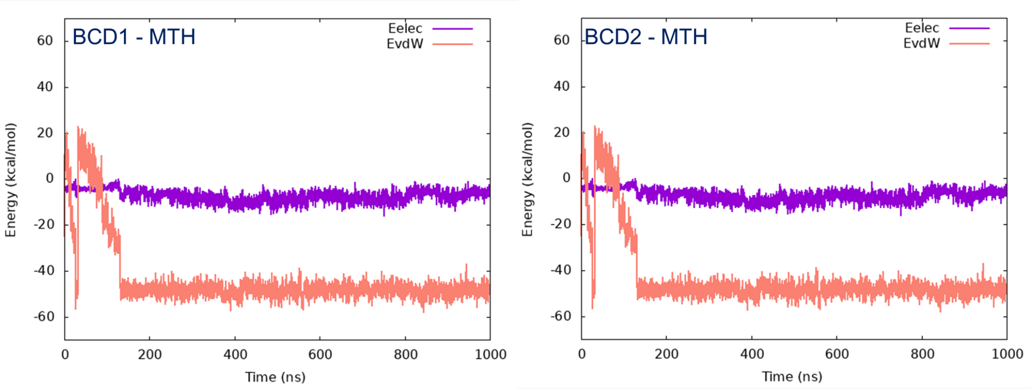


Figure S12. Time evolution of the LIE between the m-THPC and the BCD units for the second replica.

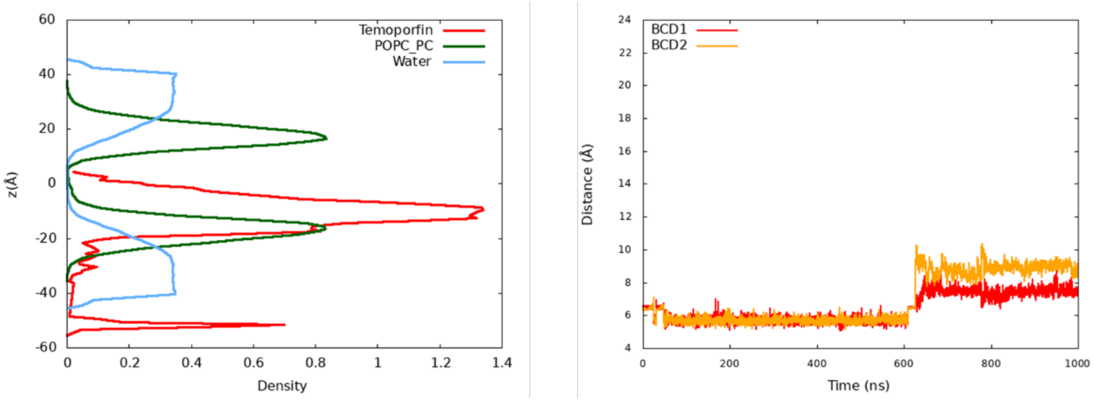


Figure S13. Density Profile (left) and time evaluation of COM distances between each cyclodextrin units and m-THPC (right) for Replica-3.

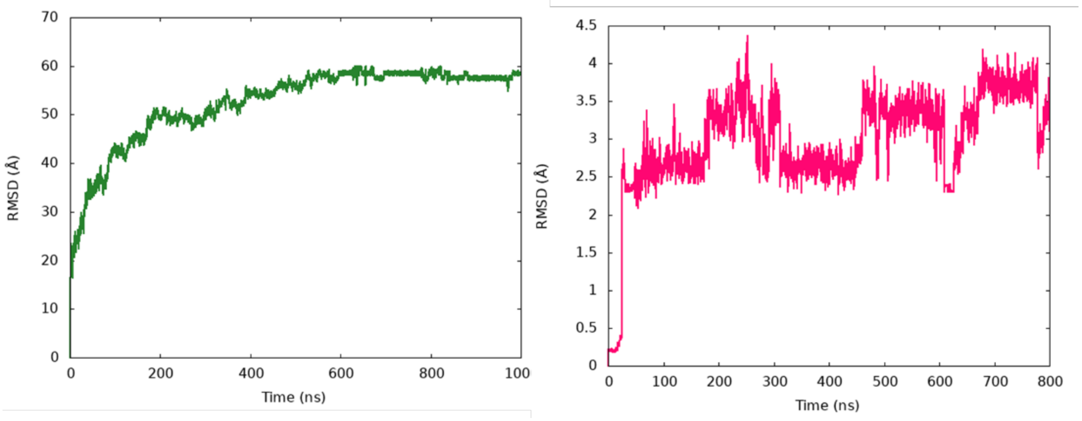


Figure S14. Root mean square deviation plots during 1μs equilibration MD of Replica-1 of the drug delivering complex in presence of the lipid bilayer. A) RMSD for the lipid and B. the complex.

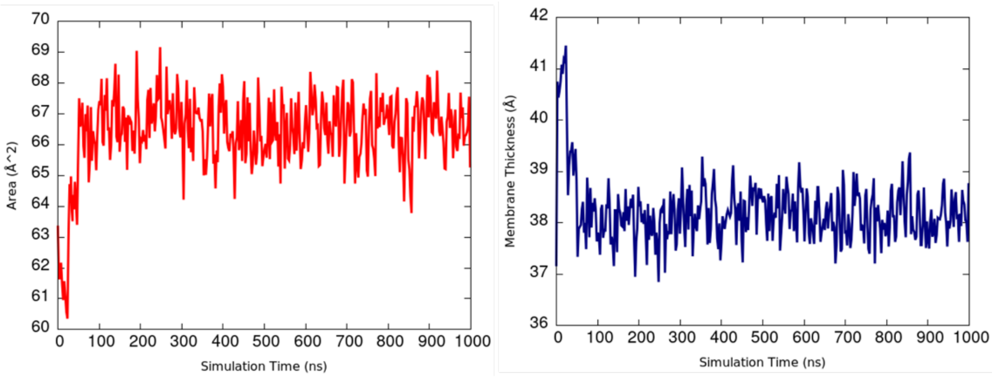


Figure S15. Time series of the area per lipid (red, left) and membrane thickness (navy blue, right) for Replica-3

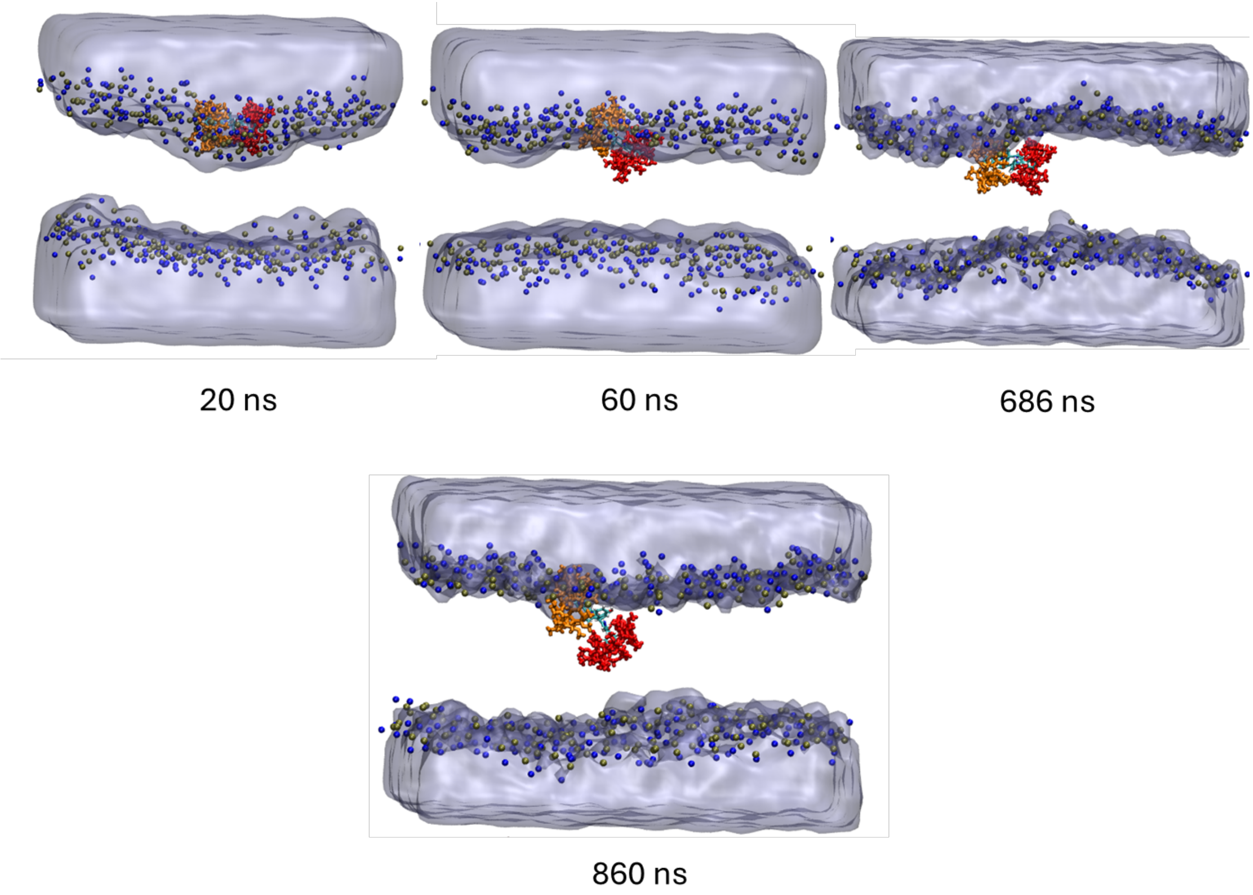


Figure S16. Representative snapshots showing the complex internalization and its partial dissociation during Replica-3.

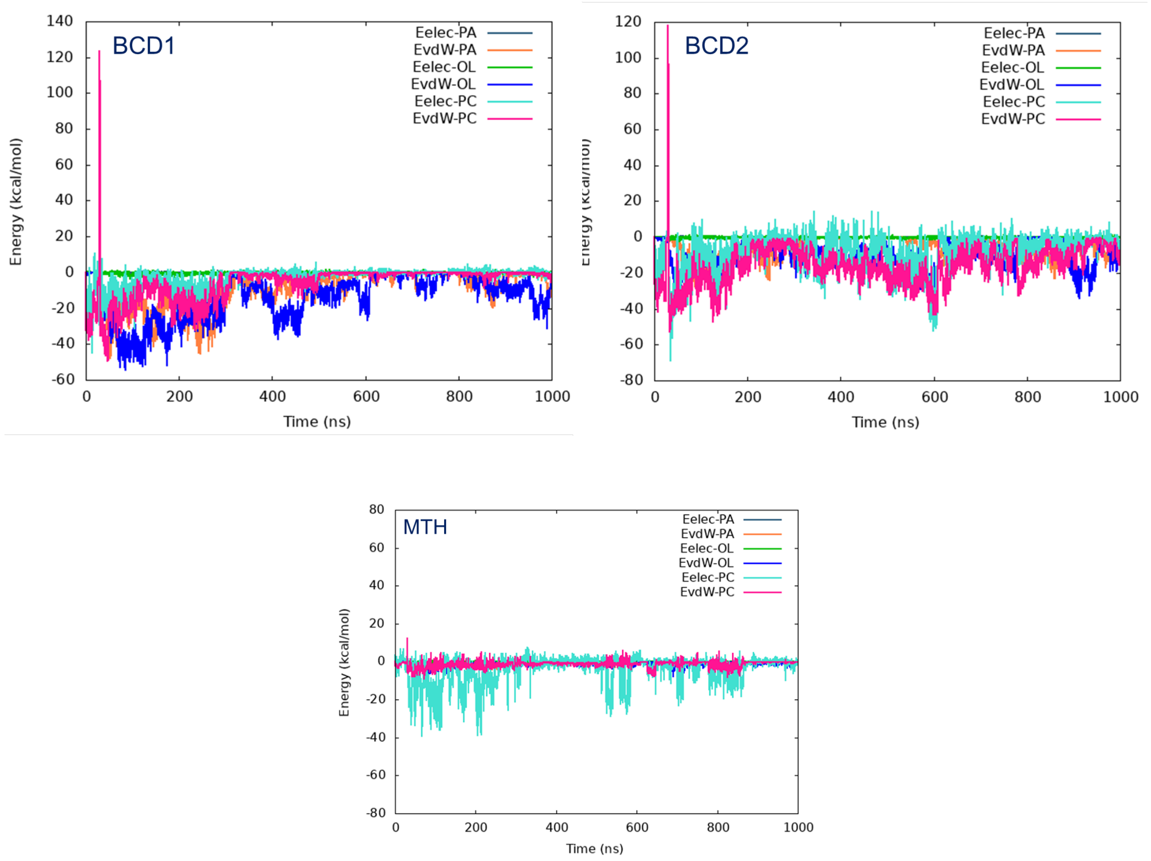


Figure S17. Time evolution of the LIE between the BCD and MTH and the lipid components for the third replica.

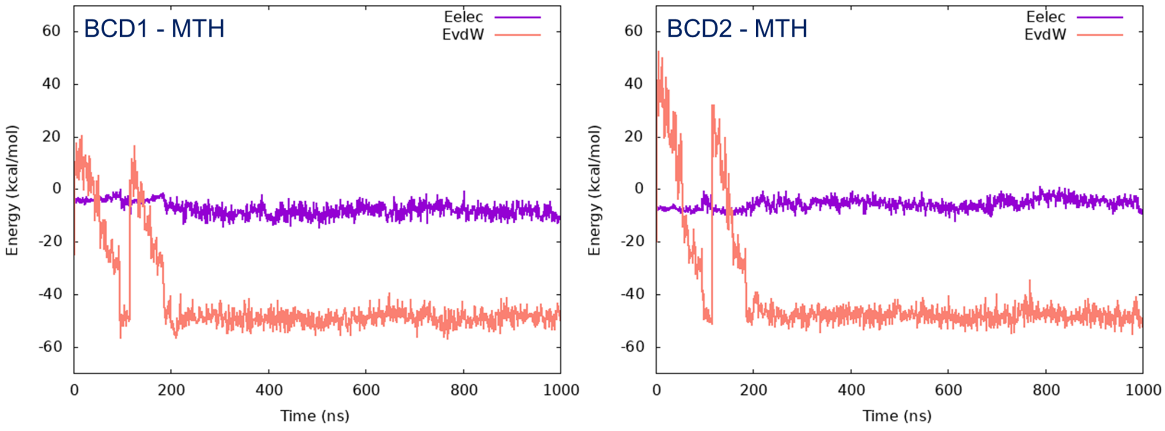


Figure S18. Time evolution of the LIE between the m-THPC and the BCD units for the third replica.
